# In *Vivo* Pharmacokinetic Study of Remdesivir Dry Powder for Inhalation in Hamsters

**DOI:** 10.1101/2020.12.22.424071

**Authors:** Sawittree Sahakijpijarn, Chaeho Moon, Zachary N. Warnken, Esther Y. Maier, Jennie E. DeVore, Dale J. Christensen, John J. Koleng, Robert O. Williams

## Abstract

Remdesivir dry powder for inhalation was previously developed using thin film freezing (TFF). A single-dose 24-hour pharmacokinetic study in hamsters, a small animal model for SARS-CoV-2, demonstrated that pulmonary delivery of TFF remdesivir can achieve plasma remdesivir and GS-441524 levels higher than the reported EC_50_s of both remdesivir and GS-441524 (in human epithelial cells) over 20 hours. The half-life of GS-4412524 following dry powder insufflation was about 7 hours, suggesting the dosing regimen would be twice daily administration. Although the remdesivir-Captisol^®^ (80/20 w/w) formulation showed faster and greater absorption of remdesivir and GS-4412524 in the lung, remdesivir-leucine (80/20 w/w) exhibited a greater C_max_ with shorter T_max_ and lower AUC of GS-441524, indicating lower total drug exposure is required to achieve a high effective concentration against SAR-CoV-2. In conclusion, remdesivir dry powder for inhalation would be a promising alternative dosage form for the treatment of COVID-19 disease.

## Introduction

The coronavirus disease 2019 (COVID-19) worldwide pandemic that is caused by Severe Acute Respiratory Syndrome-CoV (SARS-CoV-2) has strained global health care systems. Although most COVID-19 patients experienced only mild respiratory symptoms, the infection can develop into acute respiratory distress syndrome (ARDS), pneumonia and even multi-organ dysfunction which can be lethal [1]. As of December 2020, it has resulted in more than 77 million infected cases and 1.7 million deaths across the world [2]. Several therapeutic agents have been investigated for the treatment of COVID-19 such as remdesivir, favipiravir, lopinavir/ritonavir, darunavir/cobicistat, mesilate/nafamostat, chloroquine/ hydroxychloroquine, camostat, tocilizumab, eculizumab, colchicine, baricitinib, aviptadil [3] and niclosamide [4]; however, only remdesivir is currently approved by the US Food and Drug Administration for use in patients for the treatment of COVID-19 requiring hospitalization [5].

The investigational antiviral drug remdesivir (GS-5734) was originally developed for the treatment of Ebola virus infection by Gilead Sciences Inc [6]. Remdesivir is a prodrug of the parent adenosine analog, GS-441524. The MaGuigan prodrug moieties including phenol and L-alanine ethylbutyl ester help to increase lipophilicity and enhance cell permeability of the anionic phosphate moiety on remdesivir [7]. These prodrug moieties are intracellularly metabolized by esterase to an alanine metabolite (GS-704277), and further metabolized by phosphoramidase to the monophosphorylated nucleotide [8]. In the meantime, the parent nucleoside analogue core of remdesivir (GS-441524) can also diffuse into cells, and subsequently is converted to the monophosphorylated nucleotide [9]. Ultimately, the monophosphorylated nucleotide is metabolized into an active nucleoside triphosphate (NTP; GS-443902) by the host [10]. This active NTP inhibits viral RNA synthesis by competing with adenosine triphosphate (ATP) for incorporation into the nascent RNA strand [11].

Remdesivir has been repurposed for the treatment of COVID-19 as it is effective against COVID-19 in the human airway epithelial cell [12]. Recently, several on-going clinical trials have investigated the efficacy and safety of remdesivir [13–15]. In a double-blind clinical trial, severe COVID-19 patients receiving a 10-day course of remdesivir had a shorted recovery time compared to the placebo group (11 days vs. 15 days) [16]. In another clinical trial, the efficacy of a 5-day course of remdesivir or a 10-day course of remdesivir were compared with standard care in hospitalized patients with confirmed SAR-CoV-2 and moderate COVID-19 pneumonia among 105 hospitals in the United states, Europe, and Asia. The results showed the 5-day course had better clinical status distribution, compared to standard care [17].

Currently, remdesivir is only available as a lyophilized powder for reconstitution and intravenous infusion and concentrate solution for dilution and intravenous infusion [6]. Although the plasma concentration of remdesivir following IV administration can achieve two out of four reported IC50 values and two out of three reported IC90 values for the prodrug [18], the current dosage forms are limited to only hospitalized patients, excluding outpatient care. Therefore, alternative dosage forms of remdesivir for different routes of administration are necessary to improve the accessibility of the drug for patients besides those which are critically ill.

Pulmonary administration of remdesivir is a promising strategy to improve the efficacy of the treatment towards SAR-CoV2 infection as it can maximize localized lung tissue concentrations while avoiding systemic toxicities. Our group previously developed remdesivir dry powders for inhalation by thin film freezing [19]. Remdesivir combined with excipients (e.g., Captisol^®^, mannitol, lactose, leucine) at 80/20 (w/w) ratio showed optimal aerosol performance for pulmonary delivery [19]. Additionally, an *in vivo* pharmacokinetic study demonstrated that remdesivir-Captisol (80/20) exhibited a faster absorption of remdesivir into the systemic circulation, resulting in a higher amount of remdesivir in plasma and lower amount of GS-441524 in both plasma and lung tissue, compared to a remdesivir-leucine (80/20) formulation.

To further understand the pharmacokinetic profile of remdesivir and GS-441524 following administration by dry powder inhalation, this study aimed to determine its PK parameters from both plasma and lung tissue in healthy hamsters, a small animal model for SARS-CoV-2.

## Materials and Methods

### Chemical and reagents

Remdesivir for sample preparation was purchased from Medkoo Biosciences (Research Triangle Park, NC, USA). Remdesivir, GS-441524, and its heavy isotope internal standards were purchased from Alsachim (Illkirch-Graffenstaden, France). Leucine was purchased from Fisher Scientific (Pittsburgh, PA, USA). Sulfobutylether-beta-cyclodextrin (SBECD, Captisol^®^) was kindly provided by Ligand Pharmaceuticals, Inc. (San Diego, CA).

### Preparation of dry powder for inhalation

Two formulations were prepared for the pharmacokinetic study including remdesivir-Captisol (REM-CAP; 80/20 w/w), and remdesivir-leucine (REM-LEU; 80/20 w/w). Remdesivir and excipients (i.e., Captisol^®^ and leucine) were dissolved in a mixture of acetonitrile/water (50/50 *v/v)* at a 0.3% w/v solids content. The solution was dropped onto a rotating cryogenically cooled stainless-steel drum through an 18-gauge syringe needle. The drum surface temperature was controlled at approximately −100 °C. The frozen formulations were collected in a stainless-steel container filled with liquid nitrogen and then transferred into a −80 °C freezer (Thermo Fisher Scientific, Waltham, MA, USA). The formulations were dried using a lyophilizer (SP Industries Inc., Warminster, PA, USA) at 100 mTorr. The drying cycle was set at −40 °C for 20 h, and then ramped to 25 °C for 20 hours, and finally secondary drying at 25 °C for 20 hours.

### Single-dose dry powder insufflation in hamsters

This study was in compliance with the Institutional Animal Care and Use Committee (IACUC; Protocol Number AUP-2019-00254) guidelines at The University of Texas at Austin. Female Syrian hamsters (Charles River, 049LVG) 35-42-days old and weighing between 80 and 130 g (average weight of 102 g) were housed in a 12-hour light/dark cycle with access to food and water ad libitum and were subjected to one week of acclimation to the housing environment. Seventy hamsters were separated into two equal groups (REM-CAP and REM-LEU).

TFF powder formulation was passed through a No. 200 sieve (75 μm aperture) to break down large aggregates into fine particles. Precisely weighed quantities of sieved TFF powder were administered to hamsters intratracheally using a dry powder insufflator (DP-4 model, Penn-Century Inc., Philadelphia, PA, USA) connected to an air pump (AP-1 model, Penn-Century Inc., Philadelphia, PA, USA). The dose of remdesivir was targeted to be 10 mg/kg. Each hamster was briefly anesthetized with isoflurane (4% induction, 2% maintenance) and placed on its back on an intubation stand. Its upper incisors were used to secure the hamster using silk at a 45° angle, with continuous delivery of anesthesia through a nose cone. A laryngoscope was used to visualize the trachea, and the blunt metal end of the insufflator device was inserted into the trachea. The sieved TFF powder was actuated into the lung using 3 puffs of the connected pump (200 μL of air per puff). The mass of powder delivered was measured by weighing the device chamber before and after dose actuations.

Following powder administration, five hamsters from each group were harvested at each time point (15 mins, 30 mins, 1, 2, 4, 8, 24 hours). Blood was drawn via cardiac puncture and immediately transferred into a heparinized microtainer (BD, 365985, Lithium Heparin/PST™ Gel,). The blood sample was centrifuged at 10,000 rpm for 1.5 minutes, and the plasma was separated and frozen on dry ice. The hamster was carefully perfused with PBS, the lung was filled with 1 mL of PBS to collect bronchoalveolar lavage (BAL) fluid, and the lung was removed, weighed and frozen. Plasma samples, BAL and lungs were kept frozen and stored at −80 °C until analysis.

### Quantification of remdesivir and its metabolites in plasma and lung tissue

For plasma samples, remdesivir and its metabolites were extracted according to the following protocol [19]. Briefly, 100 μL of plasma was combined with 100 μL of methanol containing 100 ng/mL of the heavy labeled internal standards for remdesivir and GS-441524. The samples were mixed using a vortex mixer and then centrifuged at 12,000 rpm for 15 minutes. The supernatant was collected and placed in a 96-well plate for LC/MS/MS analysis.

For lung tissue samples, remdesivir and its metabolites were extracted according to the following protocol [19]. Briefly, lung tissue samples were added into a 2 mL tube with 3.5 g of 2.3 mm zirconia/silica beads (BioSpec Producs, Bartlesville, OK, USA), and homogenized at 4800 rpm for 20 seconds. After homogenization, 1000 μL of methanol containing 100 ng/mL internal standards for remdesivir and GS-441524 was added to the tube. The tube was then vortexed and centrifuged at 12,000 rpm for 15 minutes. The supernatant was placed in a 96-well plate for LC/MS/MS analysis. Calibration standards were prepared for plasma and lung tissue in the same protocol. Remdesivir and GS-441524 standard solutions were spiked into the blank plasma and lung tissue to obtain matrix matched calibrations. The calibration range for plasma and lung tissue was 0.1–1000 ng/ml, and 50–10,000 ng/mL, respectively. The calibration range was selected to bracket sample levels measured.

Remdesivir and GS-441524 were separated on an Agilent Poroshell column (2.1 × 50 mm, 2.7 μm) (Agilent, Santa Clara, CA, USA) using a gradient of 0 to 90.25% of acetonitrile with 0.025% trifluoroacetic acid in 5 min at a flow rate of 0.35 mL/min, and a column temperature of 40 °C. In total, 10 μL of each sample was injected for analysis with an Agilent 6470 triple quadrupole LC/MS/MS system (Agilent, Santa Clara, CA, USA).

### Pharmacokinetic analysis

Remdesivir and metabolite GS-441524 plasma and lung concentrations over time data were analyzed by non-compartmental analysis using PKSolver to obtain pharmacokinetic parameters in the hamster model after inhalation of the formulations [32]. Due to the sparse sampling requirements with this animal model and to obtain lung concentrations over time a naïve pooled-data approach was used in which the noncompartmental analysis was fit to the data as if the average of the measured concentrations from the five animals at each time point were taken from a single subject. This was based on the methods previously reported for estimating population kinetics from very small sample sizes [33, 34].

## Results

The concentrations of remdesivir and GS-441524 were determined in plasma and lung tissue from healthy hamsters treated with a single 10 mg/kg remdesivir dry powder insufflation of remdesivir/Captisol (REM-CAP) or remdesivir/leucine (REM-LEU) formulation. The resulting lung tissue concentration-versus-time curves are shown in Figure 1, while the corresponding pharmacokinetic parameters are summarized in Table 1. The C_max_ of remdesivir for REM-CAP was 8-fold higher than that of REM-LEU (75.41 ng/mg VS 8.71 ng/mg, respectively), while T_max_ of remdesivir for REM-CAP was lower compared to REM-LEU (30 mins VS 24 hour, respectively). Additionally, REM-CAP exhibited higher AUC0-24 of remdesivir in lungs than REM-LEU, indicating higher absorption of remdesivir in the lung.

**Figure 1.**
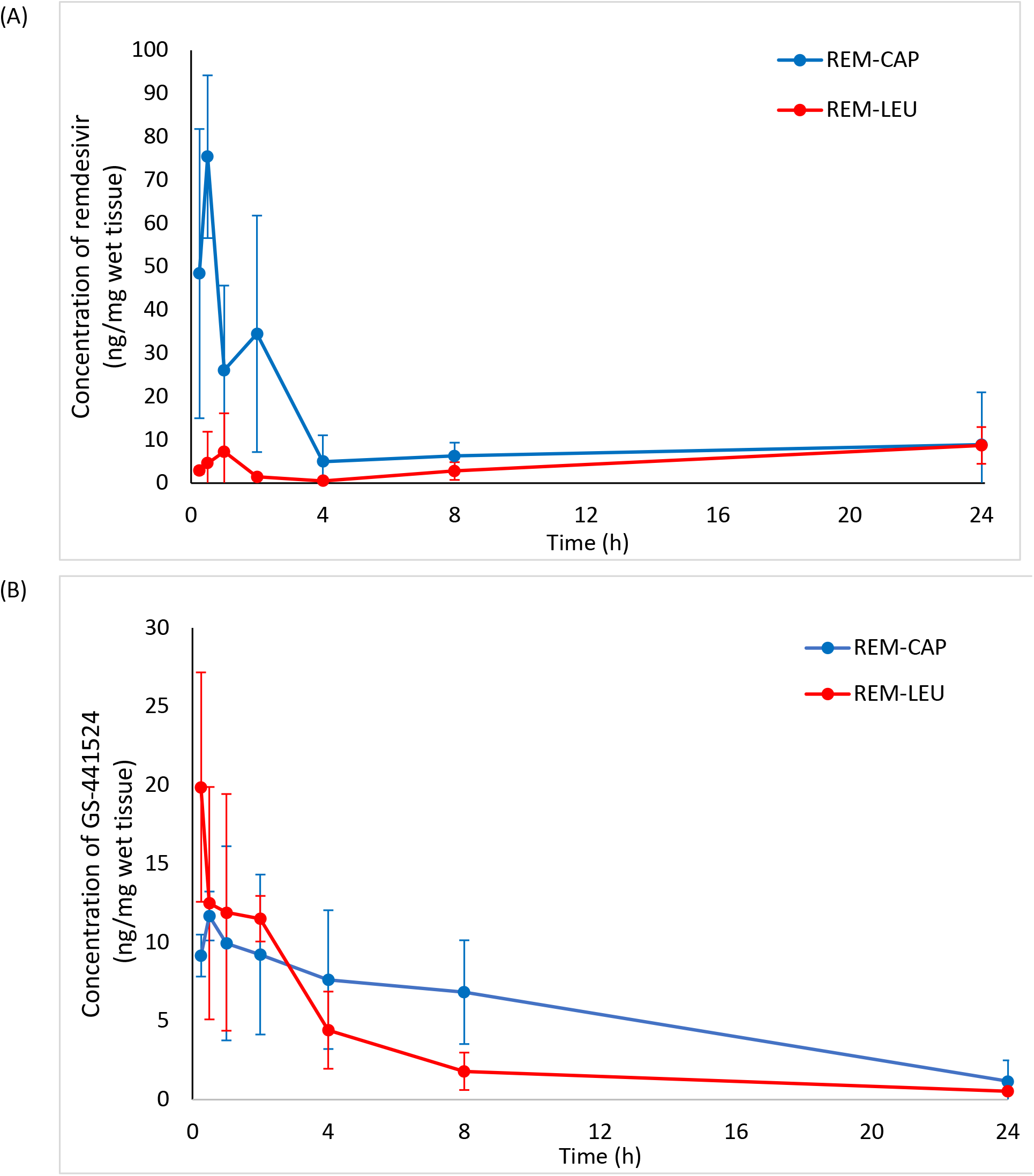
Lung concentration-time profiles of REM-CAP (remdesivir-Captisol^®^; 80/20 w/w) and REM-LEU (remdesivir-leucine; 80/20 w/w) after a single inhalation administration in hamsters; (A) remdesivir; (B) GS-441524.

**Table 1.** In vivo pharmacokinetic parameters for lung remdesivir and GS-441524 concentrations of REM-CAP and REM-LEU following a single 10 mg/kg inhalation administration.

| Pharmacokinetic parameters | REM-CAP |  | REM-LEU |  |
| --- | --- | --- | --- | --- |
|  | Remdesivir | GS-441524 | Remdesivir | GS-441524 |
| $T_{1/2}$ (h) | - | 7.38 | - | 7.12 |
| $T_{max}$ (h) | 0.5 | 0.5 | 24 | 0.25 |
| $C_{max}$ (ng/mg) | 75.41 | 11.68 | 8.71 | 19.88 |
| $AUC_{0-24h}$ (ng·h/mg) | 260.30 | 128.61 | 109.16 | 71.39 |
| $AUC_{0-inf}$ (ng·h/mg) | - | 141.11 | - | 76.85 |
| $MRT_{0-inf}$ (h) | - | 9.46 | - | 6.94 |
| $V/F$<br>(mg/kg)/(ng/mg) | - | 0.75 | - | 1.34 |
| $Cl/F$<br>((mg/kg)/(ng/mg)/h) | 0.00147 | 0.00345 | 0.00140 | 0.00209 |

For the level of metabolite in the lungs, the AUC0-24 of GS-441524 of REM-CAP was about 1.7 times higher than that of REM-LEU (128.61 ng·h/mg VS 71.39 ng·h/mg, respectively). Despite this, REM-LEU exhibited higher C_max_ of GS-441524 (19.88 ng/mg VS 11.68 ng/mg) and shorter T_max_ of GS-441524 (15 mins VS 30 mins) as compared to REM-CAP. GS-441524, in both formulations, was eliminated in a biphasic pattern with a distribution phase and an elimination phase. The half-life of GS-441524 for both formulations are similar (7.12 hours for REM-CAP and 7.38 hours for REM-LEU).

The systemic *in vivo* pharmacokinetics of drug absorption from the lungs was also investigated in hamsters. Figure 2 shows the comparison of mean remdesivir and GS-441524 plasma concentration-time profile from each formulation. The pharmacokinetic parameters following a single dose dry powder insufflation calculated using a non-compartment model are presented in Table 2. Overall, both formulations have similar plasma profiles of remdesivir and GS-441524. Rapid remdesivir concentration decay was observed in both formulations 30 mins following pulmonary administration before reaching the elimination phase. Similarly, GS-441524 plasma concentration for both formulations reached the maximum at 2 hours, and continuously decreased after 2 hours following pulmonary insufflation.

**Figure 2.**
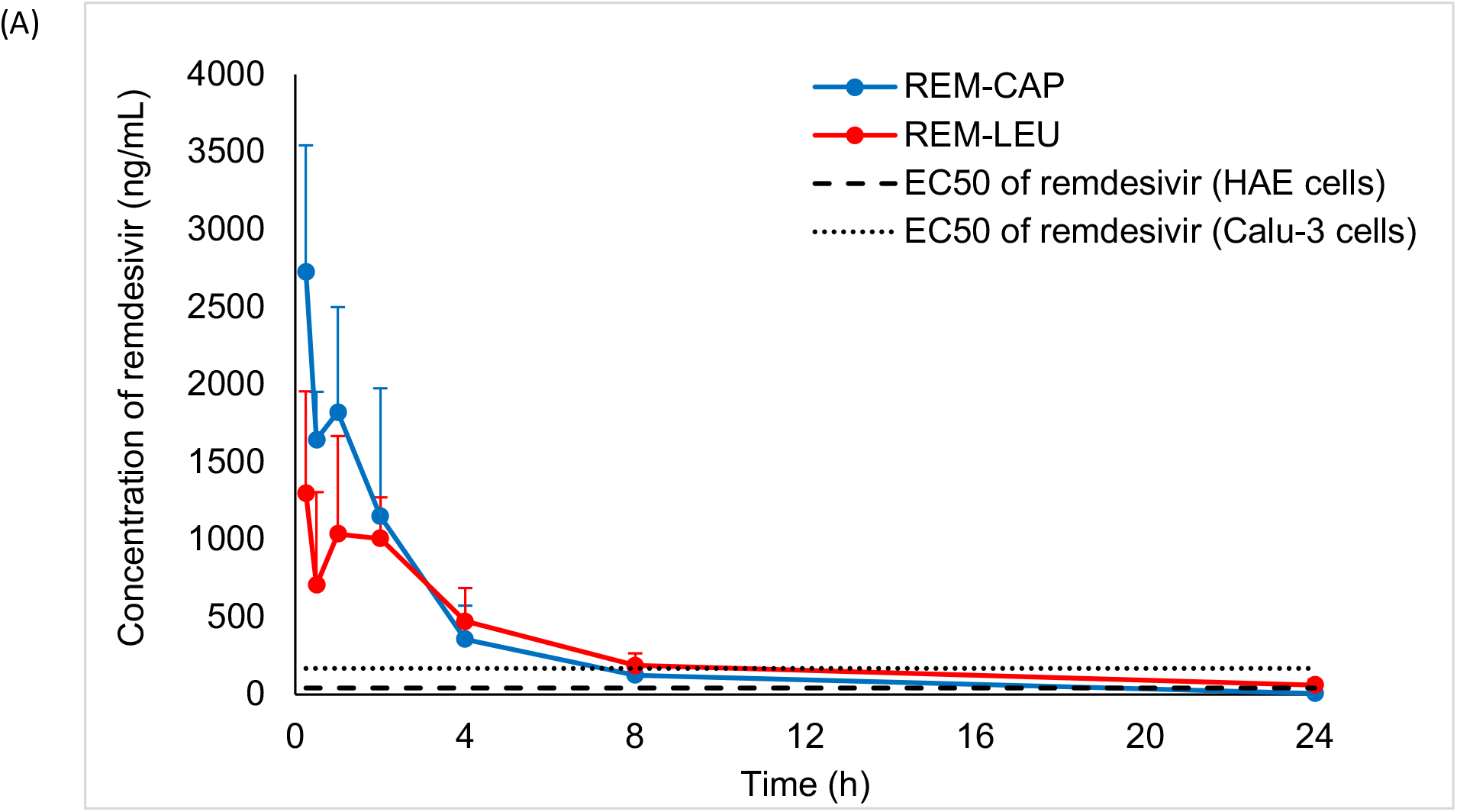
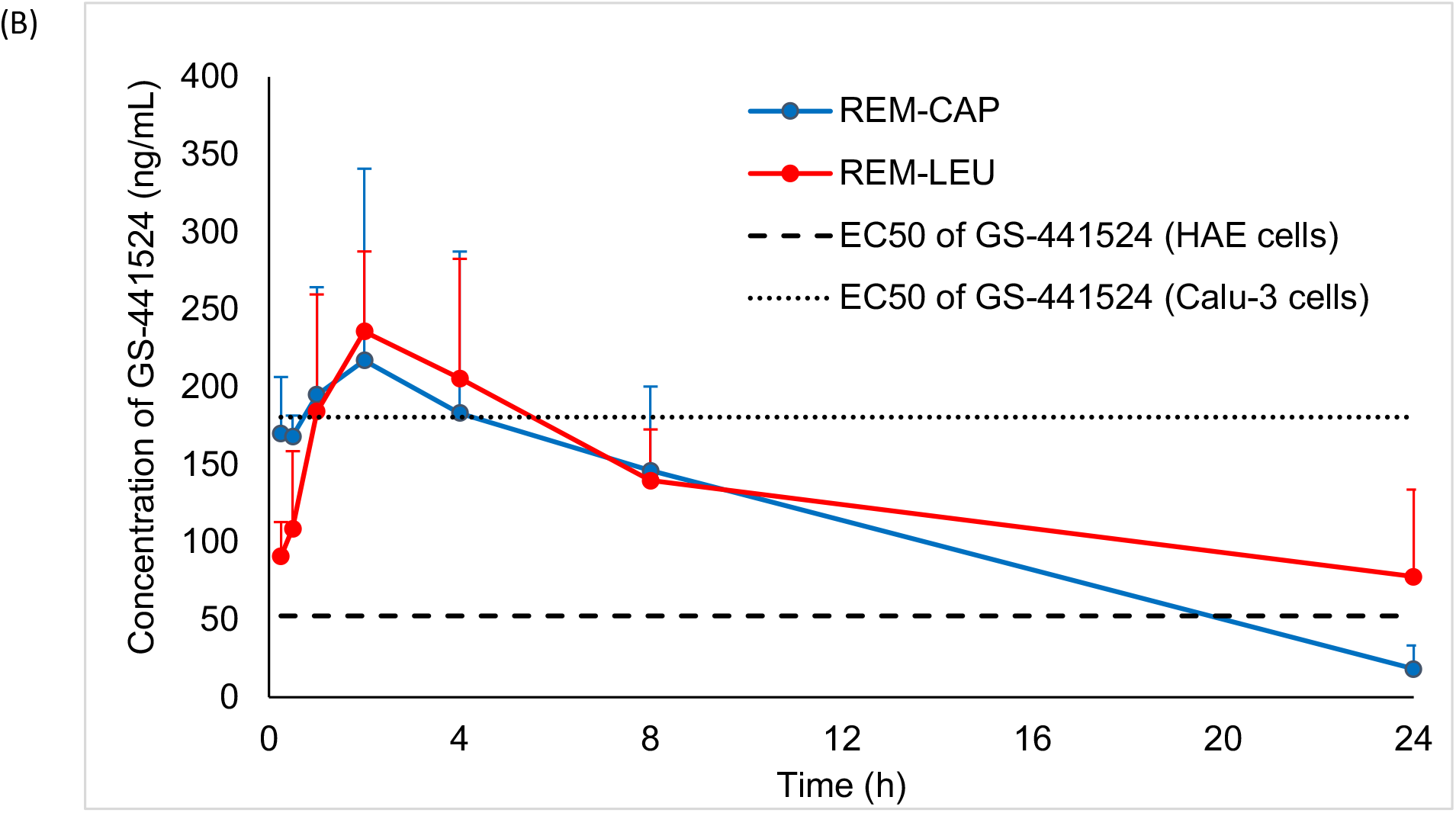
Plasma concentration-time profiles of REM-CAP (remdesivir-Captisol^®^; 80/20 w/w) and REM-LEU (remdesivir-leucine; 80/20 w/w) after a single inhalation administration in hamsters; (A) remdesivir; (B) GS-441524. Dash line and dot line represent EC_50_ of remdesivir and GS-441524 in human epithelial cells (HAE) [20], and continuous human lung epithelial cell line (Calu-3) [21], respectively.

**Table 2.** In vivo pharmacokinetic parameters for plasma remdesivir and GS-441524 concentrations of REM-CAP and REM-LEU following a single 10 mg/kg inhalation administration.

| Pharmacokinetic parameters | REM-CAP |  | REM-LEU |  |
| --- | --- | --- | --- | --- |
|  | Remdesivir | GS-441524 | Remdesivir | GS-441524 |
| $T_{1/2}$ (h) | 3.65 | 6.04 | 5.62 | 14.26 |
| $T_{max}$ (h) | 0.25 | 2 | 0.25 | 2 |
| $C_{max}$ (ng/mg) | 2726.74 | 217.10 | 1298.65 | 235.78 |
| $AUC_{0-24}$ (ng·h/mL) | 6779.83 | 2736.11 | 6647.90 | 3191.93 |
| $AUC_{inf}$ (ng·h/mL) | 6818.37 | 2895.69 | 7139.81 | 4792.48 |
| $MRT_{0-inf}$ (h) | 3.26 | 8.11 | 7.29 | 21.03 |
| $V/F$<br>(mg/kg)/(ng/mg) | 0.0077 | 0.0301 | 0.0113 | 0.0429 |
| $Cl/F$<br>((mg/kg)/(ng/mg)/h) | 0.0015 | 0.0035 | 0.0014 | 0.0021 |

Although similar absorption patterns of remdesivir and GS-441524 were observed, the pharmacokinetic parameters were slightly different. REM-CAP exhibited higher mean remdesivir plasma concentration 15 mins following pulmonary administration when compared to REM-LEU (2726.74 ng/mL VS 1298.65 ng/mL, respectively). However, both formulations have similar AUC_0-24_ and AUC_0-inf_ of remdesivir, indicating similar extent of drug absorption into the systemic circulation.

For GS-441524 plasma levels, although no significant difference in C_max_ and T_max_ of GS-441524 between two formulations was observed, the AUC_0-24_ of GS-441524 from REM-LEU was slightly higher than that of REM-CAP (3192.93 ng/mL VS 2736.11 ng/mL, respectively). Similarly, REM-LEU showed higher AUC_0-inf_ of GS-441524 compared to REM-CAP (4792.48 ng/mL VS 2895.69, respectively).

Interestingly, the half-life of remdesivir from REM-LEU was longer than that of REM-CAP (5.62 VS 3.65 hours, respectively). Likewise, REM-LEU exhibited a longer half-life of GS-441524 in plasma, compared to REM-CAP (14.26 h VS 6.04 h, respectively). This agrees with the mean residence time (MRT) of remdesivir and GS-441524. REM-LEU exhibited longer MRT_0-inf_ of remdesivir and GS-441524, indicating remdesivir and GS-441524 were cleared from the plasma more slowly than from REM-CAP formulation.

## Discussions

The *in vivo* pharmacokinetic results showed that pulmonary administration can produce plasma concentrations that achieve higher than the 50% maximal effective concentration (EC_50_). The EC_50_ is used to quantify the *in vitro* antiviral efficacy of drugs. Since the activity of antiviral agents depends on the cell type used for viral propagation, viral isolate, and viral quantification [18], several EC_50_ values of remdesivir and GS-441524 against SARS-CoV-2 in different cell lines are reported [20, 22–24]. Agostini et al. reported EC_50_ of remdesivir and GS-441524 against SAR-CoV-2 in human airway epithelial (HAE) to 70 nM (42.18 ng/mL) and 180 nM (52.43 ng/mL), respectively [20]. According to the prescribing information of Veklury^®^, the EC_50_ of remdesivir in HAE cells and continuous human lung epithelial cell line (Calu-3) is 9.9 nM (5.97 ng/mL) and 280 nM (168.73 ng/mL), respectively [21]. Pruijssers et al. also reported that the EC_50_ of remdesivir and GS-441524 in Calu-3 2B4 cells is 280 nM (168.73 ng/mL), and 620 nM (180.6 ng/mL), respectively [24]. Despite differences in the reported EC_50_ values, plasma concentrations following dry powder pulmonary insufflation of both formulations were higher than these reported EC_50_ values for remdesivir and GS-441524 in HAE cells at least over 20 hours, and higher than the remdesivir EC_50_ in Calu-3 cells for over 8 hours and likewise 4 hours for GS-441524.

The plasma GS-441524 concentration-time profiles in our hamster study are consistent with our previous pharmacokinetic study in rats. In the rat PK study, the average C_max_ of REM-CAP and REM-LEU was in the range of 220-264 ng/L. Likewise, the AUC_0-24_ and AUC_inf_ of both formulations in the rats were in the range of 2115.3-2778.5 ng·h/mL, and 2397.8 - 3204.9 ng·h/mL, respectively, which are close to the values in our hamster study. Despite the fact that different species have different drug metabolism rates, both studies indicated that pulmonary administration of remdesivir can achieve GS-441524 concentrations higher than the EC_50_.

Our *in vivo* pharmacokinetic results also provide useful information about dosing interval and dosing regimen. GS-441524 remained in the lungs for about 8 hours before plateauing (Figure 1B). The half-life of GS-441524 in the lungs of both formulations was about 7 hours. Therefore, the suggested dosing regimen of inhaled remdesivir would be twice daily.

An effect of excipients on the pharmacokinetics parameters was also observed in this study. REM-CAP exhibited a faster and greater absorption of remdesivir in the lung, as the results showed shorter T_max_, greater C_max_ and higher AUC of remdesivir in the lung. Additionally, REM-CAP produced a greater C_max_ of remdesivir in plasma when compared to REM-LEU. The hamster results agree with our previous study in rats demonstrating that the presence of Captisol^®^ resulted in the faster absorption of remdesivir compared to the leucine formulation [19]. This is possibly related to the properties of Captisol^®^. As reported in the literature, Captisol^®^ is a sulfobutyl ether derivative of β-cyclodextrin that can produce complexes with poorly soluble lipophilic drugs, and therefore enhance aqueous solubility and dissolution [25], and the bioavailability of drugs [26]. Since REM-CAP contains remdesivir and Captisol^®^ in an 80/20 weight ratio (i.e., 10:1 molar ratio of remdesivir:Captisol), approximately 10% of the remdesivir (on a weight basis) is complexed with Captisol^®^ in a solubilized form [19]. According to Eriksson et al., drug in the solution state exhibited a faster absorption than drug in dry powder form in an isolated perfused lung model since it can bypass the dissolution process which is the rate-limiting step in the absorption process when administering poorly water-soluble drug particles [27]. Another study by Tolman et al. also demonstrated that the complexation of drug and Captisol^®^ can enhance the absorption of voriconazole to systemic circulation, since AUC_plasma_ to AUC_lung_ ratio of the nebulized voriconazole formulation with Captisol^®^ was 8-fold higher than that of nanostructured amorphous voriconazole powder formulations following pulmonary administration (0.64 vs. 0.08, respectively) [28, 29]. Based on several studies, the inclusion of drug and Captisol^®^ is likely to improve the dissolution of remdesivir in lung fluid and to enhance absorption of remdesivir into the airway epithelial cell, as well as increase the absorption rate into the systemic circulation [30, 31].

Interestingly, our current study demonstrated that the complexation of Captisol^®^ and remdesivir appeared only to have an effect on the absorption rate, but not for the extent of drug absorption into the systemic circulation. The slower lung absorption of remdesivir of REM-LEU did not affect the absorption into systemic circulation as both formulations showed similar AUC_0-24h_ and T_max_ of remdesivir in plasma. Moreover, in terms of the efficacy of antiviral agents, pulmonary administration of REM-LEU can produce a higher C_max_ of GS-441524 with a shorter T_max_ and a lower AUC of GS-441524, indicating a lower total exposure of GS-441524 is needed to produce high GS-441524 levels in the lung for inhibiting virus replication. Therefore, REM-LEU would be a favorable formulation for further development.

## Conclusions

we have demonstrated that dry powder administration can deliver remdesivir to the lungs and subsequently can be converted to GS-441524 both in the lungs and plasma of hamsters. The level of remdesivir and GS-441524 in plasma was sufficiently high to provide antiviral activity. The elimination rate of remdesivir and GS-441524 from the lungs will be useful information for future efficacy studies. Lastly, dry powder remdesivir for inhalation is a promising way to improve the treatment of COVID-19 and provide an alternative dosage form on an outpatient basis.

## Acknowledgments

The authors would like to acknowledge the Drug Dynamics Institute and TherapeUTex core facility for conducting the animal study.

## References

1. Singhal, T., A Review of Coronavirus Disease-2019 (COVID-19). Indian J Pediatr, 2020. 87(4): p. 281.–286.

2. COVID-19 Dashboard by the Center for Systems Science and Engineering (CSSE) at Johns Hopkins University (JHU). 2020, Johns Hopkins University.

3. Scavone, C., et al., Current pharmacological treatments for COVID-19: What’s next? Br J Pharmacol, 2020. 177(21): p. 4813–4824.

4. Brunaugh, A.D., et al., Broad-Spectrum, Patient-Adaptable Inhaled Niclosamide-Lysozyme Particles are Efficacious Against Coronaviruses in Lethal Murine Infection Models. bioRxiv, 2020: p. 2020.09.24.310490.

5. Administration, U.S.F.D. FDA Approves First Treatment for COVID-19. October 22, 2020; October 22, 2020:[

6. Summary on compassionate use: Remdesivir Gilead. 2020, European Medicines Agency.

7. Alanazi, A.S., E. James, and Y. Mehellou, The ProTide Prodrug Technology: Where Next? ACS Medicinal Chemistry Letters, 2019. 10(1): p. 2–5.

8. Yan, V.C. and F.L. Muller, Advantages of the Parent Nucleoside GS-441524 over Remdesivir for Covid-19 Treatment. ACS Med Chem Lett, 2020. 11(7): p. 1361–1366.

9. Eastman, R.T., et al., Remdesivir: A Review of Its Discovery and Development Leading to Emergency Use Authorization for Treatment of COVID-19. ACS Cent Sci, 2020. 6(5): p. 672–683.

10. Amirian, S.E. and J.K. Levy, Current knowledge about the antivirals remdesivir (GS-5734) and GS-441524 as therapeutic options for coronaviruses. One Health, 2020. 9: p. 100128.

11. Gordon, C.J., et al., The antiviral compound remdesivir potently inhibits RNA-dependent RNA polymerase from Middle East respiratory syndrome coronavirus. J Biol Chem, 2020. 295(15): p. 4773–4779.

12. Agostini, M.L., et al., Coronavirus Susceptibility to the Antiviral Remdesivir (GS-5734) Is Mediated by the Viral Polymerase and the Proofreading Exoribonuclease. mBio, 2018. 9(2).

13. B, C., Mild/moderate 2019-nCoV remdesivir RCT - Full Text View - ClinicalTrials.gov. 2020.

14. B, C., Severe 2019-nCoV remdesivir RCT - Full Text View - ClinicalTrials.gov. 2020. 2020.

15. McCreary, E.K. and D.C. Angus, Efficacy of Remdesivir in COVID-19. JAMA, 2020. 324(11): p. 1041–1042.

16. Beigel, J.H., et al., Remdesivir for the Treatment of Covid-19 — Final Report. New England Journal of Medicine, 2020. 383(19): p. 1813–1826.

17. Spinner, C.D., et al., Effect of Remdesivir vs Standard Care on Clinical Status at 11 Days in Patients With Moderate COVID-19: A Randomized Clinical Trial. JAMA, 2020. 324(11): p. 1048–1057.

18. Rasmussen, H.B., et al., Pulmonary administration of remdesivir in the treatment of COVID-19. The AAPS Journal, 2020. 22(6): p. 1–2.

19. Sahakijpijarn, S., et al., Development of Remdesivir as a Dry Powder for Inhalation by Thin Film Freezing. Pharmaceutics, 2020. 12(11).

20. Agostini, M.L., et al., Coronavirus Susceptibility to the Antiviral Remdesivir (GS-5734) Is Mediated by the Viral Polymerase and the Proofreading Exoribonuclease. mBio, 2018. 9(2): p. e00221–18.

21. Veklury^®^ (remdesivir) for hospitalized pediatric patients [Prescribing information]. 2010, Gilead Sciences, Inc.: Foster City, CA.

22. Wang, M., et al., Remdesivir and chloroquine effectively inhibit the recently emerged novel coronavirus (2019-nCoV) in vitro. Cell Research, 2020. 30(3): p. 269–271.

23. Sheahan, T.P., et al., Broad-spectrum antiviral GS-5734 inhibits both epidemic and zoonotic coronaviruses. Science Translational Medicine, 2017. 9(396): p. eaal3653.

24. Pruijssers, A.J., et al., Remdesivir Inhibits SARS-CoV-2 in Human Lung Cells and Chimeric SARS-CoV Expressing the SARS-CoV-2 RNA Polymerase in Mice. Cell Reports, 2020. 32(3): p. 107940.

25. Lockwood, S.F., S. O’Malley, and G.L. Mosher, Improved aqueous solubility of crystalline astaxanthin (3,3’-dihydroxy-beta, beta-carotene-4,4’-dione) by Captisol (sulfobutyl ether beta-cyclodextrin). J Pharm Sci, 2003. 92(4): p. 922–6.

26. Beig, A., R. Agbaria, and A. Dahan, The use of captisol (SBE7-β-CD) in oral solubility-enabling formulations: Comparison to HPβCD and the solubility–permeability interplay. European Journal of Pharmaceutical Sciences, 2015. 77: p. 73–78.

27. Eriksson, J., et al., Pulmonary Dissolution of Poorly Soluble Compounds Studied in an ex Vivo Rat Lung Model. Mol Pharm, 2019. 16(7): p. 3053–3064.

28. Beinborn, N.A., et al., Dry powder insufflation of crystalline and amorphous voriconazole formulations produced by thin film freezing to mice. Eur J Pharm Biopharm, 2012. 81(3): p. 600–8.

29. Tolman, J.A., et al., Characterization and pharmacokinetic analysis of aerosolized aqueous voriconazole solution. Eur J Pharm Biopharm, 2009. 72(1): p. 199–205.

30. Stella, V.J. and R.A. Rajewski, Sulfobutylether-beta-cyclodextrin. Int J Pharm, 2020. 583: p. 119396.

31. Yang, K., What do we know about remdesivir drug interactions? Dlin Transl Sci, 2020. 13: p. 842–844.

32. Zhang, Y., et al., PKSolver: An add-in program for pharmacokinetic and pharmacodynamic data analysis in Microsoft Excel. Computer Methods and Programs in Biomedicine, 2010. 99(3): p. 306–314.

33. Mentré, F., et al., Sparse-Sampling Optimal Designs in Pharmacokinetics and Toxicokinetics. Drug Information Journal, 1995. 29(3): p. 997–1019.

34. Mahmood, I., Naive Pooled–Data Approach for Pharmacokinetic Studies in Pediatrics With a Very Small Sample Size. American Journal of Therapeutics, 2014. 21(4).

